# Characterizing changes in glucocorticoid receptor internalization in the fear circuit in an animal model of post traumatic stress disorder

**DOI:** 10.1101/424101

**Authors:** Emly Moulton, Marisa Chamness, Dayan Knox

## Abstract

Glucocorticoid receptors (GRs) shuttle from the cytoplasm (cy) to the nucleus (nu) when bound with glucocorticoids (i.e. GR internalization) and alter transcriptional activity. GR activation within the fear circuit has been implicated in fear memory and post traumatic stress disorder (PTSD). However, no study to date has characterized GR internalization within the fear circuit during fear memory formation or examined how traumatic stress impacts this process. To address this, we assayed cy and nu GR levels at baseline and after auditory fear conditioning (FC) in the single prolonged stress (SPS) model of PTSD. Cy and nu GRs within the medial prefrontal cortex (mPFC), dorsal hippocampus (dHipp), ventral hippocampus (vHipp), and amygdala (AMY) were assayed using western blot. The distribution of GR in the cy and nu (at baseline and after FC) was varied across individual nodes of the fear circuit. At baseline, SPS enhanced cyGRs in the dHipp, but decreased cyGRs in the AMY. FC only enhanced GR internalization in the AMY and this effect was attenuated by SPS exposure. SPS also decreased cyGRs in the dHipp after FC. The results of this study suggests that GR internalization is varied across the fear circuit, which in turn suggests GR activation is selectively regulated within individual nodes of the fear circuit. The findings also suggest that changes in GR dynamics in the dHipp and AMY modulate the enhancing effect SPS has on fear memory persistence.

## Introduction

Glucocorticoid receptors (GRs) are ligand-gated transcription factors. Upon binding with glucocorticoids they leave the cytoplasm (cy) and enter the nucleus (nu) as dimers (i.e. GR internalization) where they bind to GREs to regulate transcription (1-4). GRs can also enter the nucleus as monomers and interact with other transcription factors (e.g. AP-1) to indirectly regulate transcriptional activity (2).

Glucocorticoid release during fear conditioning (FC) has been implicated in fear memory consolidation (5-8) and specifically GR activation in the basolateral amygdala (BLA) and dorsal hippocampus (dHipp) are critical for fear memory consolidation (9-12). Changes in GR function have been consistently implicated in post traumatic stress disorder (PTSD) with an enhancement in GR levels (inferred via hormonal experiments or direct measurement on lymphocytes) being reported (13-20).

Given the role of GRs in fear memory consolidation, it is reasonable to infer that enhanced GR expression in PTSD contributes to persistent traumatic fear memory that is characteristic of PTSD (21-23). However, other studies have shown that administration of glucocorticoids shortly after trauma prevent the development of PTSD (24, 25) and can enhance the efficacy of exposure therapy in treating PTSD (26). Thus, it is currently unclear how GRs contribute to PTSD symptoms. Indeed characterization of GR internalization across the fear circuit during fear memory formation and how this process is affected by traumatic stress is lacking.

Single prolonged stress (SPS) refers to serial exposure to restraint, forced swim, and ether and is a validated animal model of PTSD (27-29). SPS exposure increases GR expression in the dHipp and mPFC (30-33) and leads to the formation of fear memory that is difficult to extinguish (i.e. persistent fear memory) (31, 32, 34-37). These two symptoms are characteristic of PTSD (18, 20, 38-40). Thus, SPS is an appropriate animal model to examine how traumatic stress might lead to changes in GR internalization in the fear circuit. In this study we used western blot to assay cy and nu GRs in the medial prefrontal cortex (mPFC), amygdala (AMY), dHipp, and ventral Hipp (vHipp) at baseline and after FC (see Figure 1).

**Figure 1.**
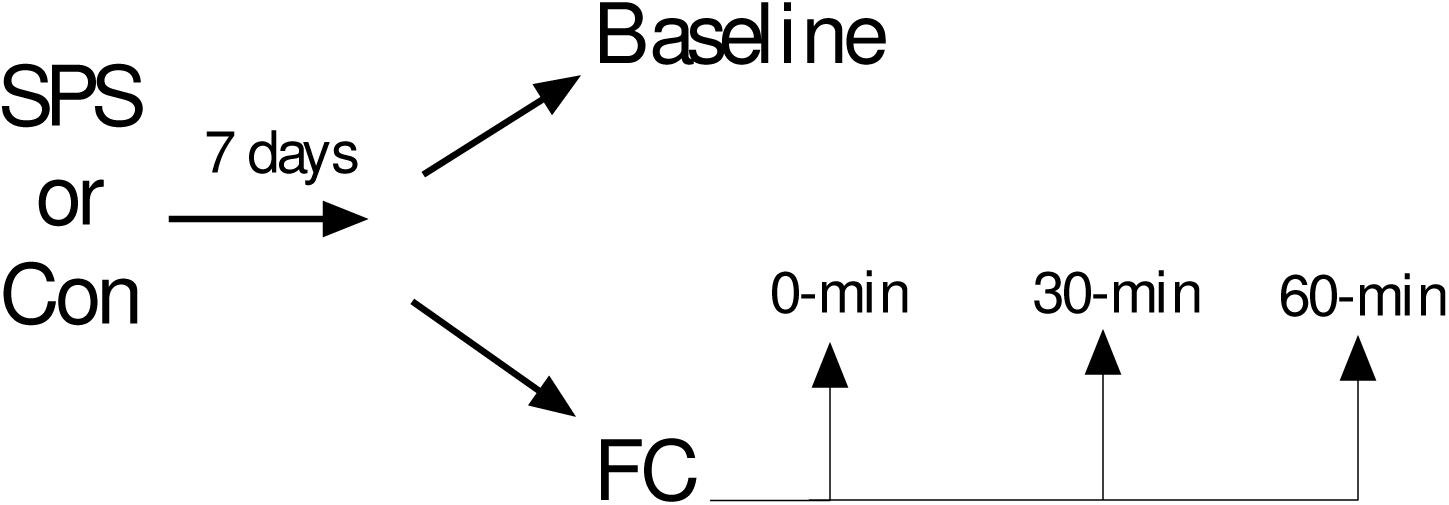
Experimental design used in this study.

These substrates were selected, because they are critical nodes of the fear circuit (41-45). Results suggest that distribution of GRs in the cy and nu at baseline and after FC was varied across these nodes of the fear circuit. The effects of SPS on cy and nu GR levels at baseline and after FC was restricted to the dHipp and AMY. SPS increased cyGR levels in the dHipp at baseline, but decreased cyGR levels in the dHipp after FC. SPS decreased cyGR levels in the AMY at baseline, but increased cyGR levels in the AMY after FC. SPS also disrupted GR internalization in the AMY brought on by FC.

## Material and Methods

### Animals

Eighty-eight male Sprague-Dawley rats (~ 150 – 250 g upon arrival) obtained from Charles River Inc. were used in this study. Upon arrival, rats were housed in pairs during a five day acclimation period with ad libitum access to food and water. Following SPS and control procedures, rats were individually housed and restricted to 23g/day of standard rat chow per the manufacturer’s recommendation (LabDiet St. Louis MO) with ad libitum access to water. Experiments commenced following the animals’ acclimation period. The rats were on a 12 hour light/dark cycle and all experimental procedures were performed in the animals’ light cycle between the hours of 9:00 am and 2:00pm. All experiments were approved by the University of Delaware Institutional Animal Care and Use committee following guidelines established by the NIH.

### SPS and Behavioral Procedures

All rats were randomly assigned to the SPS or control stress group prior to SPS. SPS was conducted as previously described (33, 46) and consisted of two hours of restraint, followed by 20 minutes of forced swim, then ether exposure until general anesthesia was induced. Control rats were placed into a novel room in their home cages while SPS occurred. A post-stress incubation period of seven days was allowed to elapse prior to experimental testing, because this is necessary to observe SPS effects (32, 33).

SPS and control rats were randomly assigned to one of four groups: baseline, FC0, FC30, or FC60. Rats in the baseline treatment were removed from the housing colony and immediately euthanized in order to determine baseline GR levels. All other rats were removed from the housing colony and subjected to FC. FC sessions were conducted as previously described (32, 34) using six MedAssociates (Fairfax VT) operant boxes. Briefly, FC consisted of five presentations of a 10 second auditory conditioned stimulus (CS, 2 kHz, 80dB) that co-terminated with a 1 second, 1mA footshock unconditioned stimulus (UCS). All FC sessions began with a 210s baseline period and had inter-stimulus intervals (ISIs) of 60s. All rats subjected to FC were removed from the operant boxes and euthanized either immediately (FC0), 30 minutes (FC30), or 60 minutes (FC60) following the cessation of FC. These time points were selected, because previous studies have shown that corticosterone levels are elevated immediately after FC, sustained for approximately 30 minutes, but after this time point begins to decrease (47-49). As a result, these time points are appropriate for examining changes in GR internalization induced by enhanced adrenal corticosterone release.

All animals were euthanized via rapid decapitation and their brains were immediately extracted and flash frozen in isopentane chilled on dry ice. Brains were then stored in a -80°C freezer until further processing. To dissect brain regions, brains were thawed to -13°C cryostat (Leica CM1350) and 300 μm coronal sections through the mPFC, AMY, dHipp, and vHipp were taken and these brain regions dissected out and placed into 1.5 mL microtubes. Dissected brain regions in microtubes were then stored in a -80°C freezer.

### Western Blot

All brain sections were treated to separate cy and nu fractions using a method described by Spencer et al., (50). We empirically tested this protocol to ensure it was successful at separating cy and nu fraction (see Appendix). Dissected brain regions were homogenized in 250 μL of buffer (50mM Tris buffer, 10% sucrose, 1mM EDTA, 0.5mM DTT, 1mM benzamidine, 0.3mM PMS Fl) by a motor-driven homogenizer (Fisher Scientific, PowerGen125). The homogenate was then then centrifuged (2,000 x g) for five minutes at 4°C to obtain a supernatant and rough pellet. The supernatant was centrifuged at 14,800 x g for 45 minutes at 4°C and the supernatant treated as the cy fraction of brain tissue.

The rough pellet from the initial centrifuge treatment was used to obtain the nu fraction from dissected brain regions. The pellet was washed twice in 400 μL of buffer and then resuspended in 150 μL of buffer that had a high concentration of NaCl (50mM Tris buffer, 0.5M NaCl, 10% sucrose, 1mM EDTA, 0.5mM DTT, 1mM benzamidine, 0.3mM PMS Fl). Samples were placed on a fixed speed vortex mixer and the suspension was incubated in ice for 1 hour with frequent shaking. Following incubation, samples were centrifuged at 8,000 x g for 15 minutes at 4°C. The supernatants from these samples were treated as the nu fraction of brain tissue.

The protein concentration of cy and nu fractions were increased using protein concentrator columns (GE Healthcare, Vivaspin 500). Protein assay was then performed on each sample per manufacturer’s directions (Pierce BCA Protein Assay Kit). 0.5X Laemmli sample buffer was mixed with approximately 15 μg of protein from each sample. These samples were stored in a -80°C until western blot. Protein samples were heated at 70°C for 7 minutes before being loaded into 10% Tris-HCl polyacrylamide gels and separated by SDS polyacrylamide gel electrophoresis. Separated proteins were electrophoretically transferred from gels to nitrocellulose membranes. The membranes were subsequently left to dry for 30 minutes at 37°C followed by rehydration washes in 0.5 M Tris-buffered saline (TBS). Blots were blocked for 1 hour at room temperature in TBS containing 5% non-fat milk. Nitrocellulose membranes were probed for GR and β actin (reference protein) by incubating overnight at room temperature with a polyclonal rabbit GR antibody (1:50, Santa Cruz Biotechnology, M-20) and a mouse β-actin antibody (1:2000, Cell Signaling Inc. 8H10D10) in TBS. After 18-20 hours, the membranes were subjected to several washes in 0.5 M TBS with 0.1% Tween-20 then a 3 hour incubation at room temperature with polyclonal goat anti-rabbit (800CW) (1:500, Li-COR) and anti-mouse IgG (680RD)
(1:5000, Li-COR) secondary antibodies in 0.5M TBS containing 0.1% Tween and 5% non-fat milk. Nitrocellulose membranes were then washed in TBS and scanned in the Li-cor Odyssey Clx scanner under the following settings: resolution – 169 μm, quality – lowest, focus offset – 0.0 mm.

### Data and Statistical Analysis

Freezing behavior was analyzed using ANY-maze (Stoelting Inc.) as previously described (32). Fear-conditioned freezing was averaged in trials that consisted of a CS and respective ISI (e.g. CS1 and ISI1) and analyzed using a stress (SPS vs. control) × trial (baseline, trials 1–5) factor design, with trial being a repeated measure. The condition factor was pooled because rats in the FC0, FC30, and FC60 levels were treated in an identical manner during FC.

In order to reduce variability in western blot data, representative rats from each independent factor (i.e. stress and condition) were always included in each protein assay and western blot. The integrated intensity (I.I.) of GR and β-actin protein bands were scored using ImageStudio. The profile curves of all bands were inspected to ensure that there was significant I.I. signal above the background within each lane in every western. For all statistical analyses cyGR and nuGR were expressed relative to cyβ-actin and nuβ-actin to yield relative GR levels. We also divided relative nuGR levels (nuGR/nuAct) into relative cyGR levels (cyGR/cyAct) to yield a single measure of GR internalization (i.e. nGR/cGR) in all brain regions.

A stress x fraction (cy vs. nu) factor design was used to examine the effect of SPS on relative cyGR and nuGR levels at baseline, with fraction being a repeated measure. T-test (SPS vs. control) was used to analyze baseline nGR/cGR ratios. A stress x fraction x condition (FC0, FC30, FC60) factor design was used to examine changes in relative cyGR and nuGR levels after FC. A stress x condition factor design was used to examine changes in nGR/cGR ratios. To specifically examine how cyGR and nuGR levels changed after FC, relative cyGR, nuGR, and nGR/cGR ratios in the condition factor were expressed as a percent change from baseline. These normalized values were subjected to separate stress x condition factor designs.

Main and simple effects were analyzed using analysis of variance (ANOVA), while main and simple comparisons were analyzed using independent, paired sample, or one sample t-test with a Bonferroni correction applied where appropriate. The reference value for all one sample t-tests was set to 100. Statistical significance was assumed with p < .05 for all statistical tests.

## Results

### Behavior

There was a main effect of trial [F_(5,125)_ = 127.615, p < .001], which suggested all rats acquired fear memory. There was also a significant stress x trial interaction [F_(5,125)_ = 2.669, p = .036]. This was driven by enhanced freezing in SPS rats during FC trial 1 of FC in comparison to control rats [t_(25)_ = 2.268, p = .032]. However, at the end of FC all animals had equivalent levels of freezing (p > .05), which suggests SPS did not alter acquisition of FC; a finding consistent with previous studies (31, 32, 34). These results are illustrated in Figure 2.

**Figure 2.**
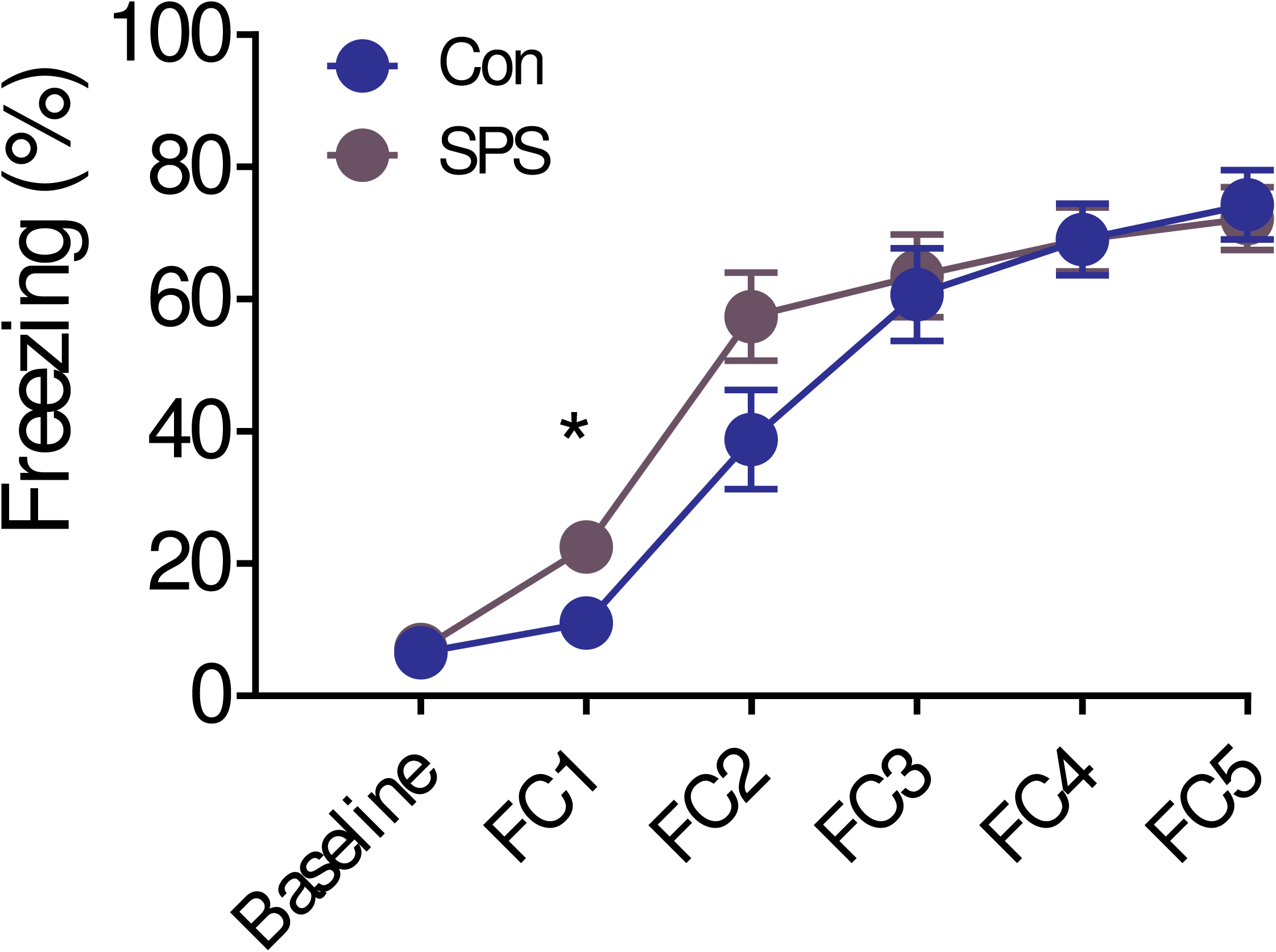
Effect of SPS on acquisition of fear conditioning (FC). Even though SPS enhanced conditioned freezing during trial 1 of FC, SPS did not affect acquisition of FC. SPS = 15, control = 12.

### Western Blot

#### vHipp

Sample western is shown in Figure 3A.

**Figure 3.**
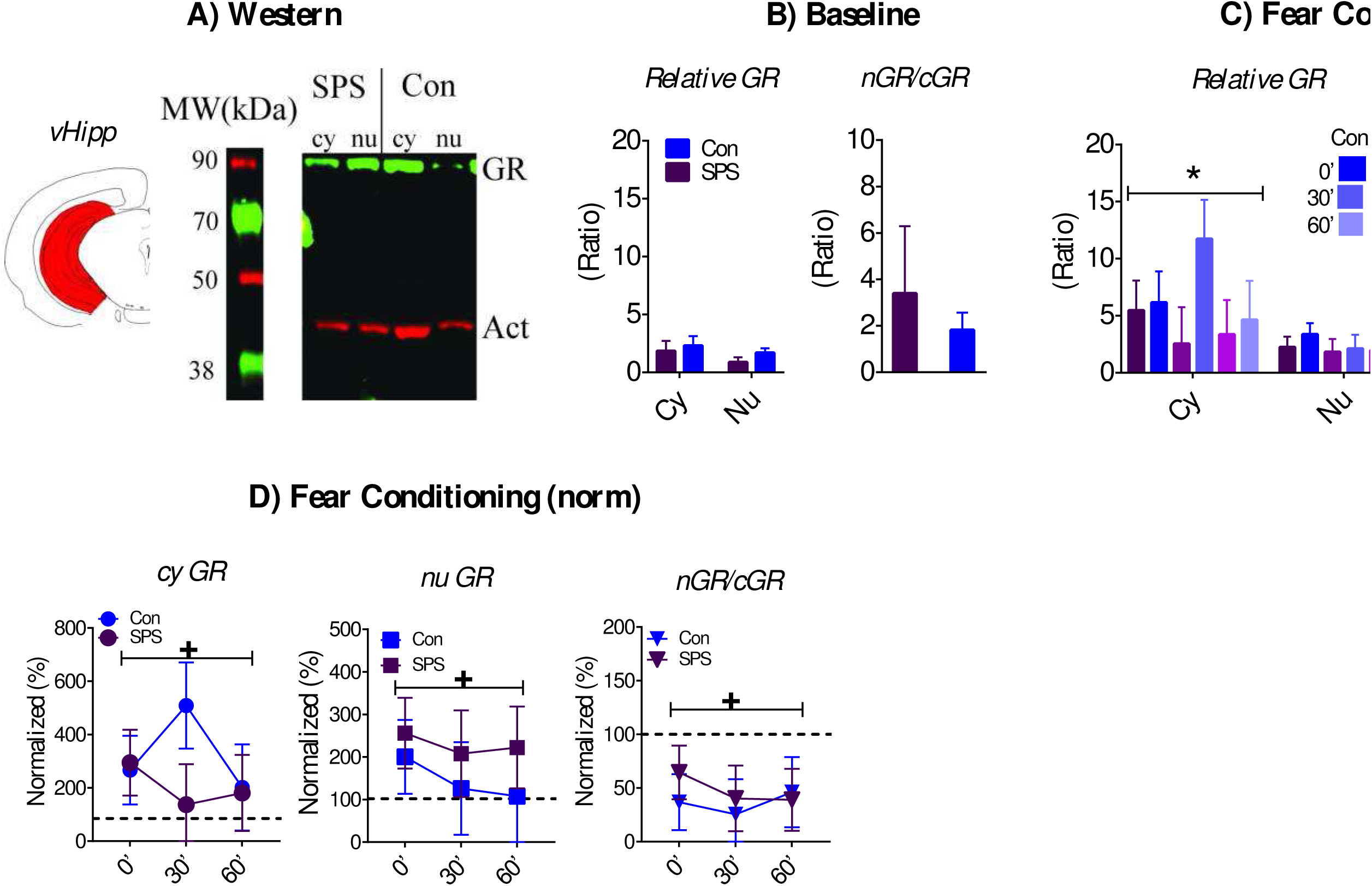
Effect of SPS on GR internalization in the vHipp. A) Cartoon of dissected brain region and representative western of cytoplasmic (cy) and nuclear (nu) vHipp samples. B) SPS had no effect on baseline GR levels or nGR/cGR ratios (SPS = 12, Con = 13). C) Cy GR levels were enhanced during FC. SPS had no effects on relative GR levels or nGR/cGR ratios after FC. D) SPS had no effect on normalized GR levels or nGR/cGR ratios, but both cy and nu GR levels were enhanced relative to baseline levels. Also, nGR/cGR levels after FC were decreased relative to baseline. + - significant one-sample t-test. SPS/0’ = 12, SPS/30’ = 8, SPS/60’ = 9; Con/0’ = 11, Con/30’ = 7, Con/60’ = 7.

Relative baseline GR levels and nGR/cGR ratios were unaffected by stress and relative baseline cyGR and nuGR levels were equivalent (Figure 3B, ps > .05). However, there was a rise in cyGR levels, relative to nuGR levels, after FC. This was revealed by a main effect of fraction [F_(1,48)_ = 11.149, p = .001; Figure 3C]. There were no stress or condition effects on relative GR levels or nGR/cGR ratios after FC (Figure 3C; ps > .05). There was no effect of stress on normalized GR levels after FC (p > .05). However, there was an enhancement in both cy and nu GR levels brought on by FC across all time points (i.e. 0’, 30’, 60’). This was revealed by significant one-sample t-test [cyGR - t_(53)_ = 2.831, p = .014; nuGR - t_(53)_ = 2.525, p = .03]. Normalized nGR/cGR ratios were unaffected by stress (p > .05), but were significantly lower after FC in comparison to baseline [t_(53)_ = 4.976, p < .001]. This suggests GR internalization in the vHipp was decreased after FC. These results are illustrated in Figure 3D.

#### dHipp

A sample western is shown in Figure 4A.

**Figure 4.**
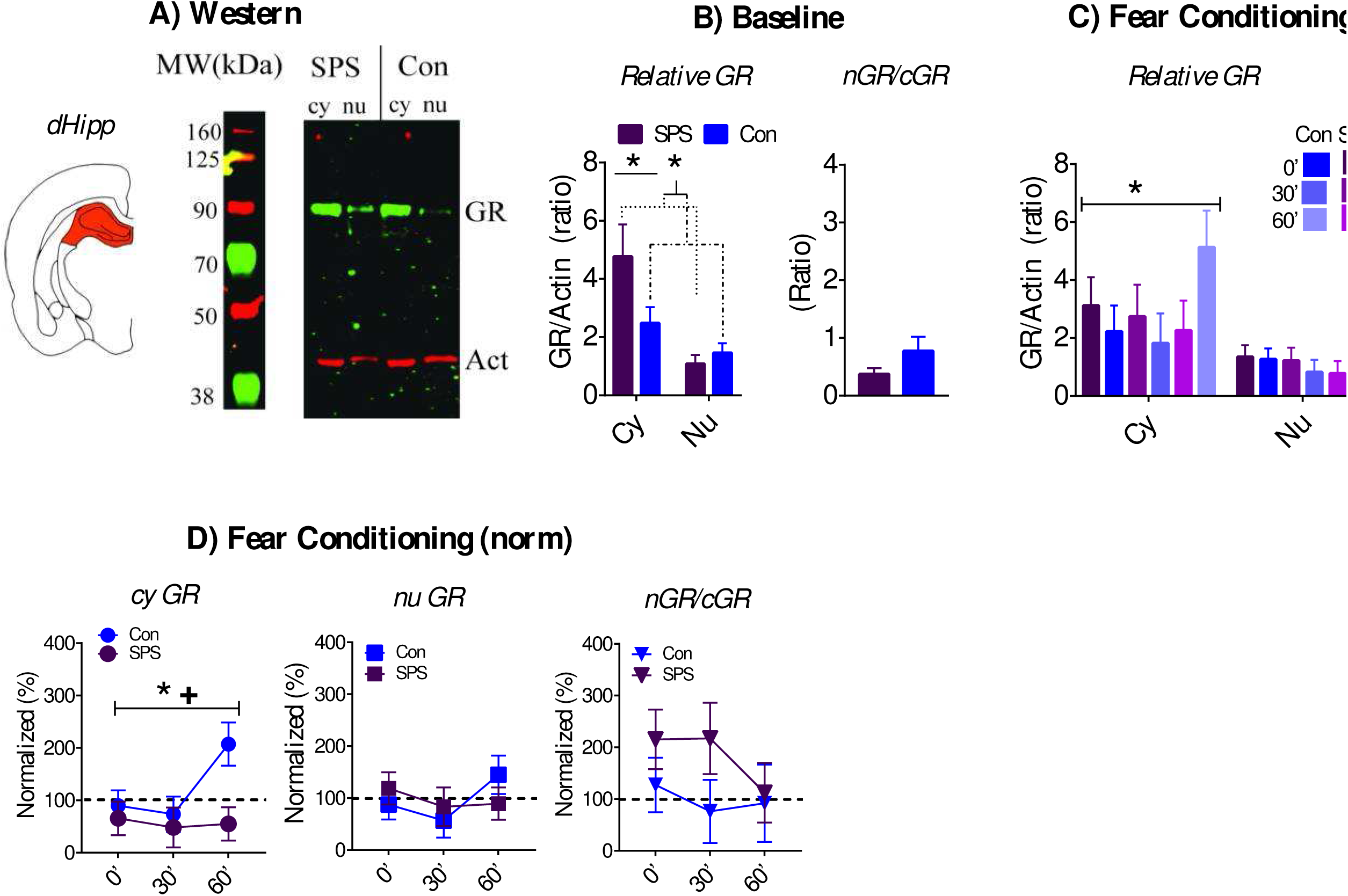
Effect of SPS on GR internalization in the dHipp. A) Cartoon of dissected brain region and representative western of cy and nu dHipp samples. B) At baseline, cy GR levels were enhanced relative to nu GR levels and this enhancement was further increased in SPS rats. SPS had no effect on nGR/cGR ratios (SPS = 9, Con = 13). C) Cy GR levels, relative to nu GR levels, were enhanced after FC. SPS had no effects on relative GR levels or nGR/cGR ratios after FC. D) Normalized cy GR levels in SPS rats were enhanced relative to control rats. This effect was most pronounced at the FC-60 time point. One sample t-test revealed that SPS decreased normalized cy GR levels after FC. SPS had no effect on normalized nu GR levels or nGR/cGR ratio. ∗ - significant effect of stress. + - significant one-sample t-test. SPS/0’ = 10, SPS/30’ = 8, SPS/60’ = 9; Con/0’ = 12, Con/30’ = 9, Con/60’ = 6.

There was a main effect of fraction [F_(1,20)_ = 57.529, p < .001] for relative GR at baseline. This reflected enhanced cyGR, relative to nuGR, in the dHipp. There was also a significant stress x fraction interaction [F_(1,20)_ = 7.345, p = .013], which was driven by SPS enhancement of cyGR relative to nuGR. This assertion was supported by significant t-test when comparing difference scores between relative cy and nu GR levels (i.e. cyGR – nuGR) for SPS vs. control rats [t_(20)_ = 2.71, p = .013]. Independent t-test for baseline nGR/cGR ratios was not significant (ps > .05). These results are illustrated in Figure 4B.

Relative cyGR was higher when compared to nuGR [main effect of fraction: F_(1,48)_ = 27.056, p < .001] after FC. There were no significant effects of stress and/or condition (ps > .05). There were no stress and/or condition effects for nGR/cGR ratios after FC (ps > .05). These results are illustrated in Figure 4C. There was a main effect of stress for normalized cyGR levels [F_(1,48)_ = 5.707, p = .021] that was driven by lower levels of cyGR in SPS rats. This effect was pronounced at the FC60 time point. Significant one sample t-test for SPS rats [t_(26)_ = 3.624, p = .001], but not control rats (p > .05), also supported the assertion that cyGR levels were decreased in SPS rats after FC. There were no significant stress and/or condition effects on relative nuGR levels or nGR/cGR ratios after FC (ps > .05). These results are illustrated in Figure 4D.

#### AMY

A sample western is shown in Figure 5A.

**Figure 5.**
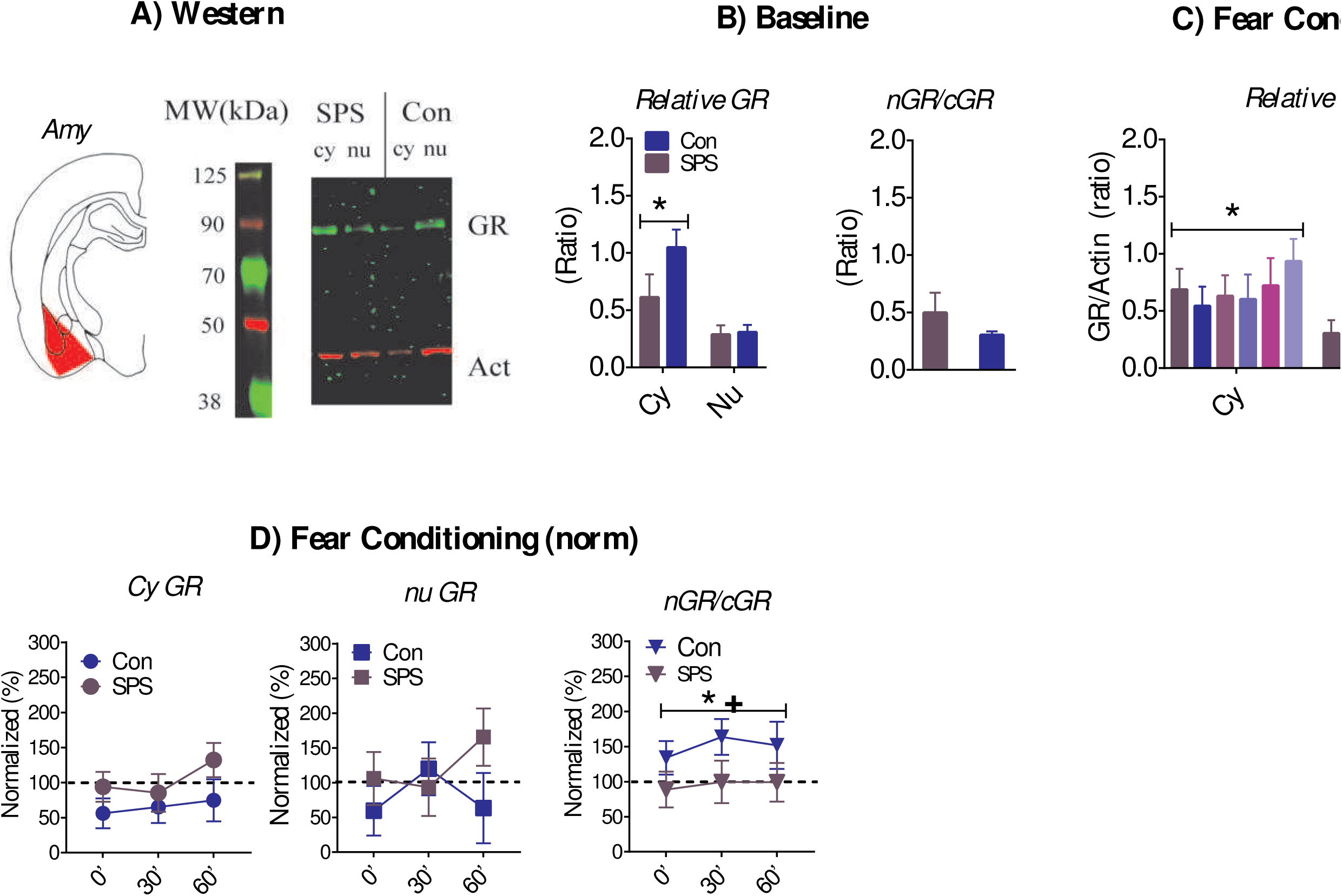
Effect of SPS on GR internalization in the AMY. A) Cartoon of dissected brain region and representative western of cy and nu AMY samples. B) Relative cy GR levels were enhanced in comparison to nu GR levels. This effect was decreased in SPS rats. This effect approached statistical significance (p = .057). SPS had no effect on nGR/cGR ratios at baseline (SPS = 6, control = 10). C) Cy GR levels were enhanced during FC. SPS had no effects on relative GR levels or nGR/cGR ratios after FC. D) SPS enhanced normalized cy GR levels. This effect approached statistical significance (p = .064). Enhanced nGR/cGR ratios after FC were observed in control rats, which suggests enhanced GR internalization. This effect was not observed in SPS rats. + - significant one-sample t-test. ^*^ significant effect of stress. SPS/0’ = 7, SPS/30’ = 5, SPS/60’ = 6; Con/0’ = 8, Con/30’ = 7, Con/60’ = 4.

There was a significant main effect of fraction [F_(1,14)_ = 28.224, p < .001] for baseline relative GR, which reflected enhanced cyGR levels relative to nuGR. This effect was attenuated in SPS rats, which was suggested by a stress x fraction interaction that approached significance [F_(1,14)_ = 4.307, p = .057]. There was no effect of stress on baseline nGR/cGR ratios (p > .05). These results are illustrated in Figure 5B.

CyGR levels were enhanced, relative to nuGR, after FC in all rats. This revealed by a significant main effect of fraction [F_(1,31)_ = 48.734, p < .001]. There were no effects of stress and/or condition on relative GRs or nGR/cGR ratios after FC (ps > .05). Relative to baseline, cyGR increased after FC in SPS rats, but not control rats. This was suggested by a stress effect that approached statistical significance [F_(1,32)_ = 3.672, p = .064]. These results are illustrated in Figure 5C. There were no effects of stress and/or condition on normalized cy and nu GR levels (ps > .05). There was a main effect of stress on normalized nGR/cGR ratios [F_(1,31)_ = 5.607, p = .024], which was driven by a failure to enhance nGR/cGR ratios after FC in SPS rats. This interpretation was supported by one sample t-test that was significant for control rats [t_(18)_ = 2.637, p = .034], but not SPS rats (p > .05). These findings suggests that FC enhanced GR internalization in the AMY and this effect was attenuated by SPS (see Figure 5D).

#### mPFC

A sample western is shown in Figure 6A.

**Figure 6.**
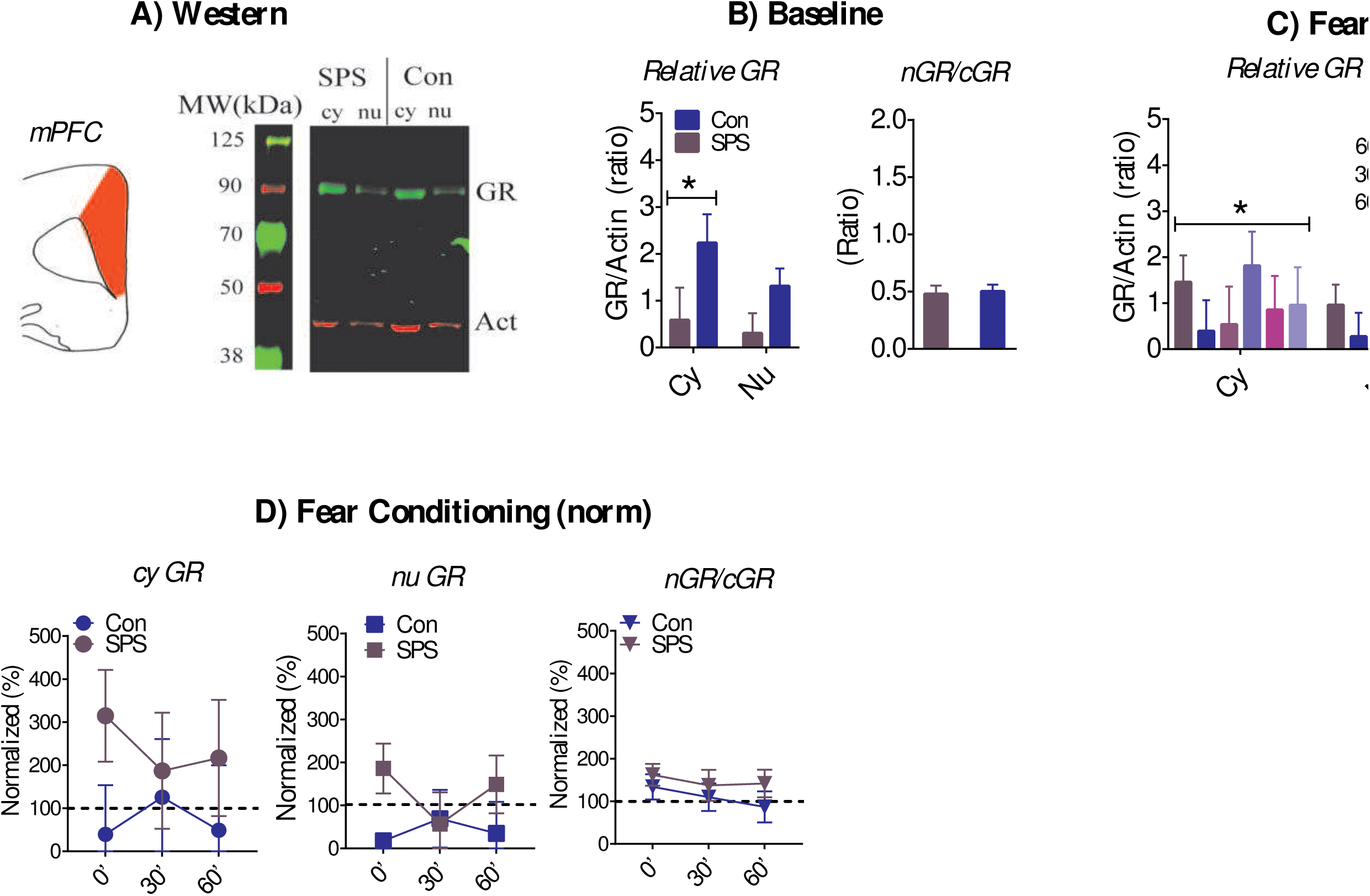
Effect of SPS on GR internalization in the mPFC. A) Cartoon of dissected brain region and representative western of cy and nu mPFC samples. B) SPS had no effect on baseline GR levels or nGR/cGR ratios, but cy GR levels were enhanced relative to nu GR levels (SPS = 7, control = 9). C) Relative cy GR levels were enhanced during FC. SPS had no effects on relative GR levels or nGR/cGR ratios after FC. D) SPS had no effect on normalized GR levels or nGR/cGR ratios. ∗ significant effect of stress. SPS/0’ = 8, SPS/30’ = 4, SPS/60’ = 5; Con/0’ = 6, Con/30’ = 5, Con/60’ = 4.

There was a main effect of fraction [F_(1,14)_ = 10.147, p = .007] for relative baseline GR, which reflected enhanced cyGR, relative to nuGR, in the mPFC. There was no significant stress effect for relative baseline GR levels or nGR/cGR ratios (Figure 6B; ps > .05). Relative cyGR levels after FC was enhanced in comparison to nuGR levels. This was revealed by a main effect of fraction [F_(1,26)_ = 4.367, p = .047].

There were no stress and/or condition effects on relative GR levels or nGR/cGR ratios after FC (Figure 6C; ps > .05). There was no effect of stress and/or condition on normalized GR levels or nGR/cGR ratios after FC (Figure 6D; ps > .05).

## Discussion

By examining changes in GR levels in the cy and nu at baseline and after FC in different neural substrates that comprise the fear circuit we were able to examine how the distribution of GRs in the cy and nu (i.e. GR dynamics) changes with SPS exposure and after FC. The results suggest there is selective regulation of GR dynamics within individual neural substrates of the fear circuit at baseline and with FC. Baseline cyGR was enhanced relative to nuGR in all brain regions except for the vHipp. FC had no effect on GR dynamics in the mPFC and dHipp, but increased cy and nu GR levels in the vHipp. In spite of this there was an overall decrease in vHipp GR internalization after FC. Enhanced GR internalization after FC was only observed in the AMY. Thus, glucocorticoid release (whether at baseline or stress-induced) does not uniformly determine GR trafficking between the cy and nu within the fear circuit.

SPS disrupted GR dynamics in the dHipp and AMY at baseline and after FC, with cyGRs being sensitive to SPS in both brain regions. SPS enhanced cyGRs in the dHipp, but decreased cyGRs in the AMY at baseline. These effects were inverted after FC with lower cyGRs in the dHipp of SPS rats, but enhanced cyGRs in the AMY. The enhancement in GR internalization in the AMY observed in control rats was disrupted by SPS. A previous study has observed that systematically inhibiting GR activation during FC exacerbates persistent fear memory induced by SPS exposure without having any effect on non-stressed rats (35). Inhibiting GR activation during FC may further inhibit GR internalization in AMY cells and decrease cyGR activation in dHipp cells. Via these processes, inhibiting GR activation during FC may enhance persistent fear memory in the SPS model. In turn, this suggests that the changes in GR function brought on by SPS can be adaptive, where GR activation during FC inhibits the development of fear memory persistence in the SPS model.

Previous studies have also shown that GR activation in the dHipp and AMY enhance memory consolidation in non-stressed rats (see Introduction), which at first appears contrary to the hypothesis that
GR activation during FC inhibits persistent fear memory in the SPS model. One explanation of this apparent discrepancy is that SPS alters GR function in the dHipp and the AMY such that activation of GRs induce different cellular effects in SPS rats when compared to non-stressed rats. Alternatively, a decrease in GR activation tends to disrupt stress adaptation (5). By decreasing GR internalization in the AMY and availability of cyGRs in the dHipp after FC, SPS may prolong the stress of FC, which renders fear memory more resistant to the inhibitory effects of extinction. Indeed previous studies have observed that the stress of FC inhibits the formation of extinction memory (51). Further research is needed to examine these possibilities.

### Substrate specific regulation of GR dynamics in the fear circuit

How might substrate specific regulation of GR dynamics occur in the fear circuit when the ligand that activates GRs originates from a single source outside of the central nervous system (i.e. adrenal cortex)? 11β-hydroxysteroid dehydrogenase types 1 and 2 (11β-HSD1, 11β-HSD2) are enzymes capable of either converting inert 11-keto forms of glucocorticoids (e.g. 11 dehydrocorticosterone) into active glucocorticoid (11β-HSD1) or metabolizing glucocorticoids (11β-HSD2). Via these mechanisms substrate specific levels of glucocorticoids can be achieved within the brain (52, 53). Both enzymes have selective expression in the brain, with high levels of 11β-HSD1 being restricted to the neocortex, hippocampus, and hypothalamus; and moderate levels of 11β-HSD2 being expressed in selective neurons in the nucleus of the solitary tract (54-57). Interestingly, genetic deletion of 11β-HSD1 results in stress resiliency (58).

GRS are phosphorylated at various sites, which alters GR function, including GR internalization (1, 59, 60). Substrate specific changes in GR phosphorylation status is observed with chronic stress and SPS (61, 62) and could be a mechanism whereby GR dynamics is selectively regulated within the fear circuit. FKBP5 is a chaperone protein for GR that inhibits GR binding by interacting with heat shock protein 90 (63-65) and has been implicated in the etiology of PTSD (47, 63, 66). These chaperone proteins have the potential to regulate GR dynamics in a substrate-specific manner by selectively lowering GR binding within neural substrates. In this study we observed rapid increases in cy and nu vHipp GRs that occurred immediately after FC and these changes may also be somewhat independent of GR internalization (see Results). Further research examining how rapid changes in GR availability might be achieved is needed, as these processes could be critical for substrate-specific regulation of GR activation in the fear circuit.

### Summary

The results of this study demonstrate that GR dynamics are varied in different neural substrates that comprise the fear circuit. This suggests that basal glucocorticoid release and stress-enhanced adrenal glucocorticoid release can have varied effects on the fear circuit via local regulation of GR activation. Furthermore, the effect of traumatic stress on GR dynamics at baseline and during fear memory formation are restricted to specific nodes within the fear circuit. Previous studies have shown that glucocorticoid administration shortly after trauma (24, 25) and during exposure therapy (26) can prevent and treat the development of PTSD. It is very likely that these treatments do not have homogenous effects on GR dynamics in the fear circuit. Characterizing how these treatments change GR dynamics at baseline and during emotional memory phenomena (e.g. FC, fear extinction) in animals models of PTSD is needed to better understand how they work and implement them in the treatment of PTSD.

## Acknowledgments

We would like to thank all of the undergraduate students who helped run behavior for this experiment.

